# Genetic potential for disease resistance in a critically endangered frog decimated by chytridiomycosis

**DOI:** 10.1101/247999

**Authors:** Tiffany A. Kosch, Catarina N. S. Silva, Laura A. Brannelly, Alexandra A. Roberts, Quintin Lau, Lee Berger, Lee F. Skerratt

## Abstract

Southern corroboree frogs (*Pseudophryne corroboree*) have been driven to functional extinction in the wild after the emergence of the amphibian fungal pathogen *Batrachochytrium dendrobatidis* (*Bd*) in southeastern Australia in the 1980s. This species is currently maintained in a captive assurance colony and is managed to preserve the genetic diversity of the founding populations. However, it is unlikely that self-sustaining wild populations can be re-established unless *Bd* resistance increases. We performed a *Bd*-challenge study to investigate the association between genetic variants of the major histocompatibility complex class IA (MHC) and genome-wide single nucleotide polymorphisms (SNPs). We also investigated differences in *Bd* susceptibility among individuals and populations, and the genetic diversity and population genetic structure of four natural *P. corroboree* populations. We found several MHC alleles and SNPs associated with *Bd* infection load and survival, provide evidence of significant structure among populations, and identified population-level differences in the frequency of influential variants. We also detected evidence of positive selection acting on the MHC and a subset of SNPs as well as evidence of high genetic diversity in *P. corroboree* populations. We suggest that low interbreeding rates may have contributed to the demise of this species by limiting the spread of *Bd* resistance genes. However, our findings demonstrate that despite dramatic declines there is potential to restore high levels of genetic diversity in *P. corroboree*. Additionally, we show that there are immunogenetic differences among captive southern corroboree frogs, which could be manipulated to increase disease resistance and mitigate the key threatening process, chytridiomycosis.

## 1 | INTRODUCTION

Southern corroboree frogs (*Pseudophryne corroboree*) are one of the world’s most threatened vertebrate species, with fewer than 50 individuals remaining in the wild (Hunter *et al.* 2010a; McFadden *et al.* 2013). This species has been driven to functional extinction after the emergence of the amphibian fungal pathogen *Batrachochytrium dendrobatidis* (*Bd*) in southeastern Australia in the 1980’s (Hunter *et al.* 2010b). *Bd* is known to infect at least 500 species of amphibians (Olson *et al.* 2013) and has caused dramatic declines and extinctions in the Americas and Australia (Berger *et al.* 1998; James *et al.* 2015). In Australia, *Bd* was first detected in Brisbane in 1978 and subsequently spread northwards and southwards along the east coast driving six species to extinction and at least seven others to near extinction (Scheele *et al.* 2017; Skerratt *et al.* 2016).

*P. corroboree* are dependent on captive assurance colonies for their continued survival. A breeding and reintroduction program (involving ~1000 adult frogs) is also underway to conserve this species in the wild (Hunter 2012; Lees *et al.* 2013). However, low recapture rates of released frogs suggest that they are still succumbing to *Bd* (Hunter *et al.* 2009), as has been observed with reintroduction efforts in other *Bd*-threatened frogs (Brannelly *et al.* 2015b; Garner *et al.* 2016; Hudson *et al.* 2016; McFadden *et al.* 2010). This is a common challenge for reintroduction programs where threats cannot be readily mitigated (e.g., infectious diseases or climate change). Most captive breeding programs aim to maintain genetic diversity and “freeze” the genetic structure of the wild source populations through time (Ballou & Lacy 1995; Schad 2007). They do not allow for adaptation to natural threats to occur and hence, species often remain vulnerable to the threatening processes that caused their declines (Schad 2007; Woodhams *et al.* 2011). Since the ultimate goal of captive breeding efforts is to reestablish self-sustaining wild populations, a better long term approach may be to apply genetic manipulation methods to increase resilience to the threatening process. Techniques such as marker-assisted selection, genomic selection, and transgenesis are well established in livestock, forestry, and crop improvement (Hayes *et al.* 2009; Hebard 2006; Jannink *et al.* 2010; Newhouse *et al.* 2014; Petersen 2017; Whitworth *et al.* 2016), but have yet to be applied to wildlife conservation (Scheele *et al.* 2014; Woodhams *et al.* 2011). Recent advances in molecular genetics have enabled the development of methods, such as genotyping-by-sequencing, for non-model species (Narum *et al.* 2013), which may allow genetic manipulation to be applied to wildlife for the first time.

Before genetic manipulation methods can be applied, detailed genomic studies must be performed to establish basic information on *Bd* immunity, including measuring phenotypic and genetic variance, and identifying genes associated with *Bd* resistance. The major histocompatibility complex (MHC) gene region has received the most attention in the context of *Bd* immunity, due to its critical role in initiating the adaptive immune response to pathogens in vertebrates. The MHC consists of several different classes of molecules, including MHC class IA that present peptides derived from intracellular pathogens to cytotoxic T cells, and MHC class IIB that present extracellular peptides to helper T cells (Bernatchez & Landry 2003; Janeway et al. 2005). Genetic polymorphism of the MHC peptide binding region (PBR) determines the repertoire of pathogens that individuals and populations can respond to, making it a good candidate marker for disease association studies and population viability estimates (Sommer 2005; Ujvari & Belov 2011).

MHC class IIB alleles, conformations, supertypes, and heterozygosity have been associated with *Bd* resistance (Bataille *et al.* 2015; Savage & Zamudio 2011; Savage & Zamudio 2016). *Bd* resistance is believed to be associated with PBR chemistries that increase the affinity of MHC molecules to bind *Bd* peptides. For example, *Bd*-resistant individuals have a distinct MHC class IIB conformation for the P9 PBR pocket that consists of an aromatic residue at β37, Aspβ57, Proβ56, and a hydrophobic β60 residue (Bataille et al. 2015). Although several studies have investigated the association between MHC class IIB and *Bd* resistance, the role of MHC class IA has not yet been examined. The intracellular life stages of *Bd* make it a likely target for MHC class IA presentation (Kosch et al. 2017; Richmond et al. 2009). Furthermore, evidence that southern corroboree frogs have high MHC class IA diversity and that selection is acting on this gene region suggests that it may play a role in *Bd* immunity in this species (Kosch et al. 2017).

Although we know that the MHC is important to *Bd* immunity, very little is known about the contribution of other genes to *Bd* resistance. Evidence from transcriptome and immunological studies suggests that multiple gene regions are involved in *Bd* immunity (e.g., Ellison *et al.* 2014b; Rollins-Smith *et al.* 2009; Rollins-Smith *et al.* 2006). Characterizing the genetic architecture (i.e., how many genes are involved and their effect size) of *Bd* resistance is fundamental to understanding how this trait evolves and for making predictions of the potential for populations to persist in the presence of *Bd*. One commonly used approach to identify the variants controlling phenotypic traits is genome-wide association studies (GWAS), using genome-wide single nucleotide polymorphism (SNP) data (Bush & Moore 2012; Quach & Quintana-Murci 2017). GWAS also allow genetic relatedness between individuals from natural populations to be estimated (i.e., realized genetic relatedness), which circumvents the necessity of pedigree data for estimating heritability (i.e., proportion of phenotypic variation due to genetic variation; *h*^2^) and permits the estimation of SNP effect size that is crucial for characterizing trait genetic architecture (Visscher *et al.* 2017).

Population differences in structure and genetic diversity can be also used to investigate the evolutionary potential of endangered species (Harrisson *et al.* 2014). Genetic diversity forms the basis for adaptation and is a major element for species conservation. Unexpectedly large differences of allele frequencies between populations can be indicative of natural selection (Lewontin & Krakauer 1973). Genome-wide selection scans (GWSS) allow the detection of regions putatively under selection by identifying markers excessively related with population structure and therefore, potential candidates for local adaptation (i.e., outlier markers; Luu *et al.* 2017; Oleksyk *et al.* 2010). Although GWSS methods are currently limited in their ability to detect polygenic traits under selection (Harrisson *et al.* 2014), a combination of approaches may detect subtle phenotype-genotype associations and changes in allele frequencies, which may shed light into the processes driving the evolution of *Bd* resistance.

The ability of natural populations to evolve disease resistance by directional selection is a key component to conserving species threatened by emerging infectious diseases. This process is dependent upon two factors: 1) the presence of phenotypic variation that differentially impacts survival or reproductive success, and 2) the existence of additive genetic variation for disease resistance (i.e., heritability; Allendorf *et al.* 2013). Here, we aimed to assess how genetic factors explain differences in *Bd* susceptibility in natural populations of southern corroboree frogs. We tested the following hypotheses using frogs experimentally exposed to *Bd*: (i) survival varies across infected individuals and populations; (ii) genetic variation at MHC loci and/or SNPs associates with infection load and survival; and (iii) MHC alleles and/or SNPs associated with survival show signatures of selection. Our approach is novel from that of previous studies in that this is the first genetic association study to investigate the association between chytridiomycosis resistance and MHC class IA, and the first to use a genome-wide approach to characterize the genetic architecture of this trait. Since our ultimate goal is to improve the success of the *P. corroboree* captive breeding program, we also analyzed the genetic structure and diversity of four of the founder populations of the captive assurance colony. We characterized their evolutionary potential and made recommendations on future research to improve the resistance of this species to *Bd*.

## 2 | MATERIALS AND METHODS

### 2.1 | ANIMAL HUSBANDRY

This study used *P. corroboree* (n=76) that were excess to the captive breeding program, and donated by the Amphibian Research Centre (Victoria, Australia). Frogs were collected as eggs from the wild from four separate populations (Cool Plains-C (n = 20), Jagumba-J (n = 18), Manjar-M (n = 22), Snakey Plains-S (n = 16)); approximate distances between populations ranged from 6 to 19 km (mean=12 km ± 5.25 SD) (for site map see Kosch *et al.* 2017; Fig 1) and raised in disease free conditions until the start of our experiment. Frogs were housed individually in 300 × 195 × 205 mm terraria with a damp and crumpled paper towel substrate (Earthcare^®^, ABC Tissue) at 18-20°C, and were fed *ad libitum* three times weekly with 5 – 10 mm crickets (*Acheta domestica*). They were misted twice daily for 60 s with reverse osmosis water, and temperature and humidity were monitored daily. Terraria were cleaned fortnightly by replacing the paper towel. The animals used in this experiment were part of a larger study (e.g., Brannelly *et al.* 2016b; Kosch *et al.* 2017). Animal ethics approval was granted by James Cook University for this study under application A1875.

### 2.2 | EXPERIMENTAL *BD* EXPOSURES

Corroboree frogs were allowed to acclimatize to their new environment for 7 d before the start of the experiment, when they were inoculated with a New South Wales strain of *Bd* (AbercrombieR-L.booroologensis-2009-LB1 passage number 11)(*Bd* treatment, n=76; controls, n=17). *Bd* zoospores were harvested from flooded TGhL petri plates and quantified using a haemocytometer. Animals were inoculated with 1×10^6^ zoospores by applying 3 mL of inoculum onto the venter. Animals were then placed in individual 40 mL containers for 6 h, and then returned to their individual terraria. *Bd* negative control animals were mock-inoculated using *Bd* negative TGhL petri plates (n = 17). We measured *Bd* infection load weekly until the end of the experiment (n = 103 d) by quantitative polymerase chain reaction (qPCR) analysis of skin swabs (Boyle et al. 2004) using the swabbing protocol and DNA extraction methods previously described (Brannelly et al. 2015a). Each qPCR analysis contained a positive and negative control, a singlicate series of dilution standards, and one replicate of each sample (Kriger *et al.* 2006; Skerratt *et al.* 2011). We monitored body condition throughout the experiment by measuring mass (to the nearest 0.01 g) and snout to vent length (SVL) weekly. Body condition was estimated by Log10(Mass+1)/Log10(SLV+1). Frogs were checked daily for general health and clinical signs of chytridiomycosis (Brannelly *et al.* 2015c) and were euthanized with an overdose of MS-222 in accordance with animal ethics guidelines if deemed moribund. Any animals that cleared infection and survived until the end of the experiment were returned to the Amphibian Research Centre.

Survival data between populations was analysed by Cox Regression analysis using the survival package in R (Therneau 2015; Therneau & Grambsch 2000). Infection loads were transformed by taking the Log10 of zoospore equivalents (ZE) + 1, and analysed using mixed models with nlme in R (Pinheiro *et al.* 2009). Constructed models included the explanatory (fixed) factors of week, population, week*population, and days survived. ANOVA was used to evaluate which models best fit the data.

### 2.3 | MHC CLASS I GENOTYPING

#### DNA extraction and PCR

DNA was extracted from various tissues (skin, muscle, kidney, toe clips) using an ISOLATE II Genomic DNA Kit (Bioline) following the manufacturer’s instructions. DNA concentration and quality was measured with a Nanodrop2000 (Thermo Fisher), and extracts were stored at −20°C until use. Polymerase chain reaction (PCR) amplification was performed using *P. corroboree* MHC class IA exon 2 primers (PcIAex2-2F1, PcIAex2-2R1), which amplify the hypervariable α1 peptide binding region (PBR) domain (Kosch *et al.* 2017). Initially, PCRs were performed with *Taq* DNA polymerase, but preliminary sequencing runs suggested that the DNA polymerase and reaction conditions were leading to sequencing errors (indicated by the occurrence of single bp changes not replicated across multiple sequences). Therefore we modified PCR conditions to minimize PCR errors and artefact formation for the remaining runs using previously described modifications (Babik 2010; Judo *et al.* 1998; Zylstra *et al.* 1998). Specifically, we switched to a high fidelity DNA polymerase (NEB Q5 High-Fidelity PCR Kit), decreased DNA template amount to 60 ng, increased annealing temperature to 67°C, increased elongation time to 3 min, and reduced cycle number to 25 (see Methods S2 for complete reaction details).

Resulting PCR products were separated by gel electrophoresis and bands of the correct size were excised and extracted with a FavorPrep Gel Purification Kit (Favorgen) following the bench protocol. PCR bands generated using the Q5 PCR kit were extracted with a different kit (NEB Monarch PCR & DNA Cleanup Kit) to inactivate the exonuclease included in the reaction and A-tailed before ligation (see Methods S2 for A-tailing reaction).

#### Cloning and sequencing

All PCR products were ligated with a pGEM^®^-T Easy Vector kit (Promega), and recombinant DNA was transformed into Top 10 competent *Escherichia coli.* Cells were grown on LB agar plates (with 100 µg/ml ampicillin and 20 µg/ml X-Gal) for 16 h at 37°C. We used blue-white screening to select 16 to 24 clones from each transformation and amplified them with SapphireAmp Fast PCR Master Mix (Takara) and M13 primers. Multiple independent PCRs were run for a proportion (n = 32) of the individuals to rule out single copy alleles and potential PCR artefacts (Table S1). We also used previously published sequence data available for a subset of the individuals (Kosch *et al.* 2017) to confirm genotypes.

PCR products were purified for sequencing by a clean-up reaction of 10 µl of PCR product, 1 U of Antarctic phosphatase (NEB), 1 U of exonuclease (NEB), and 2.6 µl of RNase-free water and the thermal cycler program: 37°C for 30 min, 80°C for 20 min, and 4°C for 5 min. Resulting purified PCR products were then shipped to Macrogen (Seoul, South Korea) for unidirectional Sanger sequencing.

#### MHC sequence analysis

Resulting sequences were analyzed with Geneious (v. 9.0.5) and identified as alleles if: (i) BLAST results indicated they were MHC Class IA sequences, (ii) they did not include stop codons, and (iii) they were present in more than one copy per individual and more than one independent PCR reaction. Alleles were named based upon MHC nomenclature rules described in Klein *et al.* (1993), and were assigned to supertypes to explore functional diversity. Supertype designation was performed by first aligning corroboree frog amino acid sequences with that of *Homo sapiens* (HLA-A; D32129.1). Next we extracted amino acid sequences from the 13 PBR pocket positions identified in previous studies (Lebrón *et al.* 1998; Matsumura *et al.* 1992) using R. We then characterized the 13 sites for five physiochemical descriptor variables: z1 (hydrophobicity), z2 (steric bulk), z3 (polarity), z4 and z5 (electronic effects) (Didinger *et al.* 2017; Sandberg *et al.* 1998) and performed discriminant analysis of principle components (DAPC) with R package adegenet (Jombart *et al.* 2010) to define functional genetic clusters. Alleles were assigned to clusters by a K-means clustering algorithm by selecting the model with the lowest Bayesian information criterion (BIC).

We tested for recombination in our nucleotide alignment with the genetic algorithm recombination detection (GARD) method executed on the Datamonkey server (Delport *et al.* 2010; Kosakovsky Pond *et al.* 2006). In MEGA7, we tested for evidence of positive selection with the Z-test of selection on three datasets: (i) the entire MHC class IA alignment, (ii) putative PBR pockets, and (iii) non-putative PBR pocket nucleotides using the modified Nei-Gojobori method (Jukes-Cantor) and 500 bootstrap replications (Kumar *et al.* 2016; Nei & Gojobori 1986). Positive selection at the codon level (dN/dS or ω > 1 with a posterior probability of > 0.95) was estimated with omegaMap (v 5.0) (Wilson & McVean 2006) following similar conditions to (Lau *et al.* 2016). Tajima’s D test of neutrality was executed in MEGA7. Evolutionary relationships among *P. corroboree* nucleotide sequences and other vertebrates were inferred by constructing Neighbor-Joining (NJ) phylogenetic trees in MEGA7. Evolutionary distances were computed using the Kimura 2-parameter gamma distributed method (K2+G) and tree node support was estimated via 500 bootstrap replicates (Felsenstein 1985).

We investigated population differences in MHC class IA diversity using five measures: (i) the number of unique alleles per population (A_P_); (ii) the number of alleles per individual (A_I_); (iii) mean evolutionary distance between nucleotide (D_NUC_) and amino acid (D_AA_) variants estimated with MEGA7 (Kumar *et al.* 2016) as the number of differences over all sequence pairs within each individual using a p-distance model; (iv) the total number of MHC supertypes per population (S_P_); and (v) the mean number of MHC supertypes per individual by population (S_I_). Number of alleles (A_I_) and supertypes (S_I_) per individual were summarized with a generalized linear model (GLM) in R assuming a Poisson distribution to model the count data. One-way analysis of variance (ANOVA) in R was used to compare population evolutionary distances between nucleotides (D_NUC_) and amino acids (D_AA_). Arlequin was used to estimate pairwise fixation index (F_ST_), population differentiation based on variance of allele frequencies among populations (Excoffier & Lischer 2010). The theoretical range of F_ST_ values is from 0 to 1, with 0 indicating complete panmixia and 1 indicating two isolated populations.

### 2.4 | SNP GENOTYPING AND QUALITY CONTROL

To investigate the genome-wide association with infection load and survival, all infected individuals (n = 76) were genotyped by Diversity Arrays Technology Sequencing (DArTseq, Canberra, Australia). This method uses hybridization-based sequencing technology implemented on an NGS platform to identify thousands of single nucleotide polymorphisms (SNPs) in one reaction (Cruz *et al.* 2013). Because high molecular weight DNA is necessary for DArTseq analysis, we examined our DNA samples by gel electrophoresis before shipping. A subset of samples from extracted skin (n = 23), were re-extracted from kidney tissue due to the presence of extensive nucleosome ladders or smearing. Samples were then diluted to 50 ng/µl with TE Buffer to a final volume of 15 µl for DArTseq analysis.

Sequence read quality was filtered for >10 Phred quality score and minimum pass percentage of 50. Initial SNP quality control was performed by DArTdb with >3 reads per SNP and >95% reproducibility. Further filtering (MAF of 2%, call rate 70%, duplicate removal) and data formatting was then performed with dartQC (<u>https://github.com/esteinig/dartQC</u>) resulting in 3,489 SNPs.

### 2.5 | GENOTYPE-PHENOTYPE ASSOCIATION ANALYSIS

#### Genome-wide association analyses

We applied more stringent quality control with the GenABEL ‘check.marker’ function to exclude SNPs with a call rate ≤ 95% and individual call rate ≤ 95% (Aulchenko *et al.* 2007). We also excluded two individuals in which the identity by state (IBS) was greater than 0.9. We evaluated Hardy–Weinberg equilibrium (HWE) independently for each population and removed SNPs if they failed this test (P≤ 0.001) in all four populations. After quality control, 3,245 SNPs remained for GWAS analysis. To investigate the associations between SNPs and the phenotypic traits we ran a separate GWAS for each phenotypic trait, three in total: (i) maximum infection load (log transformed), (ii) days survived (log transformed), and (iii) infection load per week. Maximum infection load and days survived were tested using mixed models in the R package GeneABEL (Aulchenko *et al.* 2007). Repeated measurements of infection load per week were rank transformed using the ‘rntransform’ function in GenABEL and analysed using the function ‘rGLS’ in the R package RepeatABEL (Rönnegård *et al.* 2016). We accounted for the effects of size in models (i) and (iii), sex in model (ii) and week in model (iii), as these factors were significantly associated with the trait. To account for multiple hypothesis testing, the p-value significance thresholds were adjusted with the Bonferonni equation using two different alpha thresholds (alpha=0.05, significant; alpha=1.0, suggestive) (Clarke *et al.* 2011). The heritability values (h^2^_SNP_) of each of the ten SNPs with the smallest P-values (i.e., top 10 SNPs) were estimated as V_SNP_/V_P_ (V_P_ is the phenotypic variance estimate for the phenotypic trait and V_SNP_=2pqa^2^, where p and q are the frequencies of the major and minor allele frequencies, respectively, and a is the additive SNP effect (Falconer *et al.* 1996). The top ten SNPs from each of the three GWAS were annotated by searching the NCBI non-redundant nucleotide database with the software package Blast2GO (Götz *et al.* 2008).

### 2.6 | POPULATION GENETIC ANALYSES

Several measures were used to estimate population genetic variation and thus assess long-term evolutionary potential of *P. corroboree* populations. Mean allelic richness (A_R_) was estimated using the R package PopGenReport (Adamack & Gruber 2014). Observed heterozygosity (H_O_), expected heterozygosity (H_E_) and inbreeding coefficient (F_IS_) were estimated using the R package diversity (Keenan *et al.* 2013). Effective population size (N_e_) was estimated using the software NeEstimator v.2 using the linkage disequilibrium method and a random mating model (Do *et al.* 2014). Expected heterozygosity (H_E_) is the best overall estimate of genetic variation, and can be compared to H_O_ to estimate inbreeding rates (i.e., H_E_ > H_O_ suggests excessive inbreeding) (Allendorf *et al.* 2013). Allelic richness (A_R_) measures allelic diversity while considering sample size. This method is more likely to detect population bottlenecks than H_E_ (Allendorf 1986). Effective population size (N_e_) indicates the rate of heterozygosity loss over time due to stochastic factors such as genetic drift (i.e., populations with smaller Ne have a greater rate of heterozygosity loss through time) (Allendorf *et al.* 2013; Kliman *et al.* 2008).

Pairwise F_ST_ values were calculated using the software GenePop on the web (Rousset 2008). An exact G-test was also calculated in GenePop (Markov-chain parameters: 10,000 dememorisation steps, 1000 batches and 10,000 iterations per batch) for each population pair using the G log likelihood ratio.

Outlier markers were identified using the PCAdapt R package (Luu *et al.* 2017) with K = 9 and min.maf = 0.01. The candidate loci were determined using Benjamini–Hochberg FDR (false discovery rate) control and the level of FDR was set to 0.01. To evaluate the genetic relationships among individuals, discriminate analysis of principal components (DAPC) was performed using adegenet package in R (Jombart 2008) for neutral and outlier SNPs. The a-score approach was used to assess the stability of the DAPC analyses (i.e. trade-off between power of discrimination and over-fitting). Across all 100 permutations the highest *a*-score was 0.655 for 3 PCs.

### 2.7 | POPULATION STRUCTURE BASED ON MHC CLASS IA AND SNP DATA

We used the program STRUCTURE 2.3.4 (Pritchard *et al.* 2000) to examine clustering of source *P. corroboree* populations based on either MHC class IA or SNP genotypes. For MHC class IA, because multiple MHC loci were amplified, we entered data recessive alleles based on the approach used for AFLP data sets (Falush *et al.* 2007). The four populations were incorporated into the admixture model. We determined the number of genetic clusters of individuals (K) using the method of Evanno *et al.* (2005) to calculate deltaK in STRUCTURE HARVESTER (Earl 2012). We tested a range of K = 1 to K = 5 with 10 replicates of each K, using 100,000 iterations following a burn-in period of 100,000 iterations.

## 3 | RESULTS

### 3.1 | SURVIVAL AND INFECTION LOAD OVER TIME

#### Survival

Five frogs in the *Bd*-inoculated group survived to the end of the experiment, four of which were from population M (18.2%), and one from population C (5.0%). All animals in the negative control group survived the experiment. Frogs became ill and were euthanized between day 21 and day 94 post inoculation. Cox proportional hazards regression indicated that population of origin had a significant impact on days survived (Figure 1) (Cox regression: 
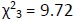
, P < 0.05), with population M surviving on average 14.2 days longer than the other three populations.

**FIGURE 1.**
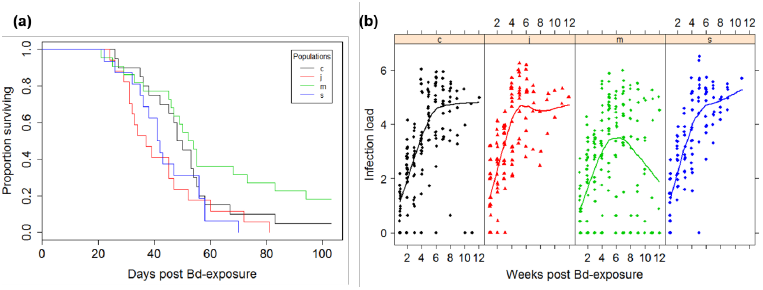
Interpopulation variation in *Bd* infection load and mortality in laboratory exposed *P. corroboree* (a) Daily survivorship in *Bd* infected frogs. (b) *Bd* infection load (log10(ZE+1)) over the course of the experiment as estimated by qPCR. Trend lines represent smooth fitted population means. In the second half of the experiment infection load trends for population M diverged from the other populations as highly infected frogs died, and the means became more influenced by survivor loads. Populations are represented by (c) Cool Plains, (j) Jagumba, (m) Manjar, and (s) Snakey Plains

#### Infection load

All *Bd*-inoculated frogs were *Bd* positive for at least two weeks during the experiment, and infection loads increased over time in all but the 5 survivors. Negative controls remained *Bd*-negative throughout the duration of the experiment. Body condition during the experiment decreased in *Bd* inoculated frogs, but not controls (Figure S1). Four frogs (5.3%; population M, n = 3; population C, n = 1) successfully cleared infection by week 12. The fifth survivor had a low infection load (6.7 ZE) at the end of the experiment and later cleared infection naturally. The overall log of infection load increased dramatically in the first half of the experiment (slope = 0.835), and then plateaued in the second half (slope = −0.225) (Figure 1). To account for this change of slope through time, the dataset was subdivided into two datasets (early < 5.5 weeks and late > 5.5 weeks) before mixed effects modelling. ANOVA results comparing models of infection load indicated that the model of best fit for both early and late datasets included days survived (transformed with a quadratic function to improve linearity), allowed infection load to vary by population, had a Week*Population interaction factor, and included individual ID as a random effect (ANOVA, AIC = 1066, χ215 = 7.321, P = 0.062; AIC = 456, 
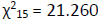
, P < 0.0001). Infection load did not differ between populations in either the early or late dataset (Mixed models, F_3,47_ = 0.507, P= 0.678; F_3,54_ = 1.540, P = 0.215), but there was a signification interaction of Population*Week in the late dataset (Mixed models, F_3,124_ = 3.156, P < 0.05) with population M having a significantly different slope than the other 3 populations (ANOVA, F_3, 185_ =14.63, P <0.001). The change in the slope of population M in the late dataset was due to the impact of the 4 surviving individuals reducing infection from moderate levels, rather than individuals recovering from high infection burdens.

### 3.2 | MHC DIVERSITY AND EVOLUTION

#### MHC allele diversity

We identified a total of 22 MHC class IA alleles, with a range of 2 to 10 alleles per individual (Table S2). Alleles blasted with high similarity to MHC IA sequences from *P. corroboree* (KX372222-KX372242, and KY072979-KY072985), *Rana clamitans* (JQ679356), and *R. temporaria* (FJ385608) (Kiemnec-Tyburczy *et al.* 2012; Kosch *et al.* 2017; Teacher *et al.* 2009). There were no differences in number of alleles per individual (A_I_) (GLM, Χ^2^ = 3.852, d.f. = 3, P = 0.278), mean evolutionary distance between nucleotide variants (D_NUC_) (GLS, F_3,71_ = 2.527, P = 0.0643), and mean evolutionary distance between amino acid variants (D_AA_) (GLS, F_3,71_ = 1.877, P = 0.141) among populations (Table 1, Figure S5). The most common MHC allele, Psco-UA*9, was present in > 70% of individuals across populations (range = 31.0% – 85.0%) (Table S3, Figure 3b). Alleles Psco-UA*24 and Psco-UA*27 were unique to population C and alleles Psco-UA*18 and Psco-UA*19 were unique to population M. FST values of the MHC class IA were significant at p < 0.05 level for three of six pairwise comparisons involving the four populations (M x C, M x J, M x S; range: 0.000 – 0.012; Table 2).

**TABLE 1.** MHC class IA genetic diversity statistics by population. (N) number of individuals, (A_P_) total number of alleles per pop, (A_I_) number of alleles per individual averaged per population, (D_NUC_) mean pairwise nucleotide diversity (p-distance), (D_AA_) mean pairwise amino acid diversity (p-distance), (S_P_) total number of supertypes per population, and (S_I_) number of supertypes per individual averaged per population. () standard deviation

|  | Population |  |  |  |
| --- | --- | --- | --- | --- |
|  | C | J | M | S |
| N | 20 | 18 | 22 | 16 |
| $A_P$ | 17 | 17 | 20 | 14 |
| $A_I$ | 5.75 (2.15) | 5.72 (1.71) | 6.41 (1.94) | 4.88 (2.28) |
| $P_A$ | 2 | 0 | 2 | 0 |
| $D_{NUC}$ | 0.187 (0.020) | 0.169 (0.023) | 0.183 (0.018) | 0.188 (0.031) |
| $D_{AA}$ | 0.322 (0.038) | 0.297 (0.034) | 0.301 (0.029) | 0.312 (0.038) |
| $S_P$ | 8 | 7 | 8 | 7 |
| $S_I$ | 4.650 (1.387) | 4.611 (1.420) | 5.409 (1.469) | 4.500 (1.789) |

**TABLE 2.**
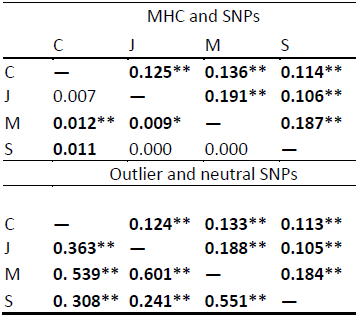
Corroboree frog population genetic differentiation (FST values). (upper graph) MHC class IA alleles (lower half) and SNPs (upper half), (lower graph) 24 outlier SNPs (lower half) and 3465 neutral SNPs (upper half). bold (p<0.05), * (p<0.01), and ** (p<0.001).

#### MHC evolution

We found no evidence of recombination between MHC alleles. There was evidence of positive selection acting on codons of the putative PBR pocket sites (dN/dS = 2.128, Z = 2.921, P ˂ 0.01), but no evidence of positive selection on the non-PBR pocket sites (dN/dS = 0.590, Z = −1.702, P = 1.000) or the entire MHC Class IA region (dN/dS = 0.693, Z = −0.027, P = 1.000). Tajima’s D value of > 0 on the entire alignment indicates that balancing selection or sudden population contraction has occurred (D = 1.09). In total, omegaMap identified 11 codons with evidence of positive selection, of which 7 sites aligned with codons of HLA-A PBR pocket positions (Figures 2 and S3; 1, 16, 26, 53, 56, 57 and 60). Three of these sites (16, 56, and 57) have been previously identified as being under positive selection in this species (Kosch et al. 2017).

**FIGURE 2.**
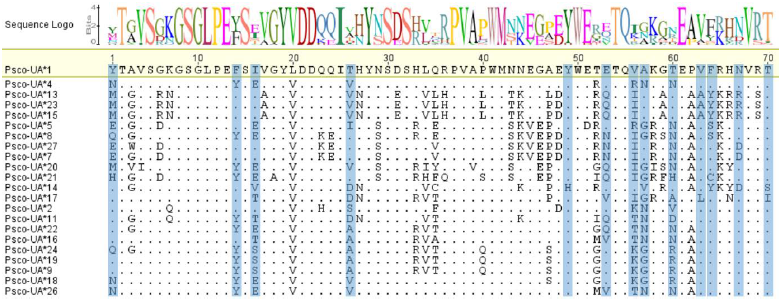
Alignment of *P. corroboree* MHC Class IA amino acid sequences. Peptide binding pocket positions from *H. sapiens* (Lebron 1998 and Matsumura 1992) are highlighted in grey. Dots indicate homology to the reference sequence Psco-UA*1

#### MHC supertype diversity

Conversion of 22 MHC alleles into functional supertypes resulted in 8 distinct supertypes, with each supertype containing one to four alleles (Figure S3). The most common MHC supertype, ST8, was present in > 80% of individuals (range = 68.0% – 90.0%) (Table S4, Figure 3a). MHC supertypes corresponded to groups of clades within the NJ phylogeny (Figure S4). Supertypes 1 and 2 were each comprised of a single allele. Supertypes 4 and 5 formed two distinct clades, while supertypes 3, 6, 7, and 8 were split into separate clades dispersed throughout the phylogeny. The number of supertypes per individual (SI) ranged from 1 to 8 (mean = 4.83 ± 1.53 SD) with no difference among populations (GLM, Χ^2^ = 2.16, d.f. = 3, P = 0.541).

**FIGURE 3.**
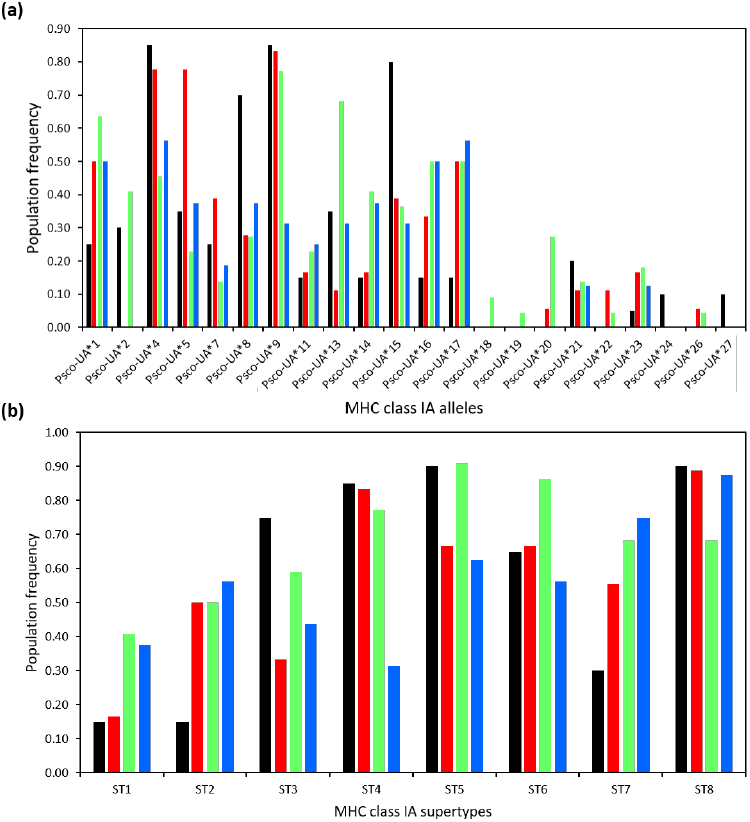
MHC class IA allele and supertype distributions among *P. corroboree* populations. (a) allele and (b) supertype (ST) frequency distribution. The incidence of *Bd* susceptibility-associated allele (Psco-UA*5) and supertype (ST8) was lowest in population M. (black) population C, (red) population J, (green) population M, and (blue) population S

### 3.3 | ASSOCIATION ANALYSIS

#### MHC association

Alleles Psco-UA*5 and Psco-UA*9 were positively associated with maximum infection load (Table S5; GLS, F_1, 74_ = 4.11, P ˂ 0.05, F_1, 74_ = 10.56, P ˂ 0.01). Allele Psco-UA*23 was negatively associated with number of days survived (Table S6; GLS, F_1, 74_ = 12.96, P ˂ 0.001). Allele Psco-UA*5 was least common in the more resistant population M (23% ± 0.19) and most common in the susceptible population J (78% ± 0.21) (Table S3, Figure 3a). Strangely, alleles Psco-UA*9 and Psco-UA*23 were relatively common in the more resistant population M (77% ± 0.19 and 18% ± 0.19 respectively). Individuals with ST8 had higher maximum infection loads than those with other STs (Table S7; GLS, F_1, 74_ = 4.49, P ˂ 0.05) and a greater chance of dying (Figure S6; GLS, F_1, 73_ = 7.29, P ˂ 0.01).

#### GWAS

The association analyses did not identify any significant SNPs after correction for multiple testing (Figures 4 and S7), although one SNP (173) was suggestively negatively associated with days survived (P=9.2e-05; Table S9; Fig 4). In general, each one of the top SNPs explained only a small proportion of the phenotypic variation. Two SNPs (1894 and 1895) were identified in the top ten markers positively associated with both maximum infection load (GenABEL) and infection load per week (RepeatABEL). BLAST results revealed that 96% of the top SNPs had sequence homologies with other amphibians, including *Xenopus tropicalis*, *X. laevis*, *Nanorana parkeri*, and *Andrias davidianus* (Table S10). Several of the top SNPs were homologous to genes that are known to impact immunity and included functions such as pathogen recognition and control and immune cell proliferation (Table 4).

**FIGURE 4.**
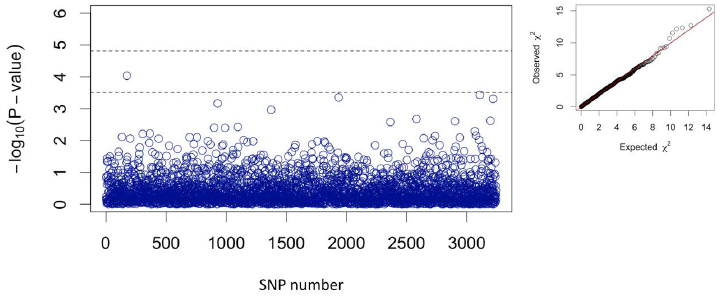
Manhattan plot with −log_10_ p-values from the association with marker genotype (SNPs) for days survived using GenABEL. The upper dashed line indicates the genome-wide Bonferroni-corrected significant threshold and the lower dashed line indicates the suggestive threshold. The QQ-plot (on the right) shows the relationship between the expected and observed distributions of SNP level test statistics

### 3.4 | GENETIC DIVERSITY AND STRUCTURE OF SNPS

PCAdapt analyses with a false discovery rate (FDR < 0.01) resulted in 3465 neutral and 24 outlier SNPs. F_ST_ values from all SNPs ranged from 0.106-0.191, F_ST_ values from neutral SNPs ranged from 0.105-0.188 and FST values from outliers ranged from 0.241-0.601 with populations M and J being the most differentiated and populations S and J being the least differentiated for all datasets (i.e. including all SNPs, neutral and outlier SNPs; Table 2). DAPC plots using neutral SNPs showed all the populations clustering independently. When using only the 24 outlier SNPs, population M was distinctively separated from the remaining populations (Figure 5).

**FIGURE 5.**
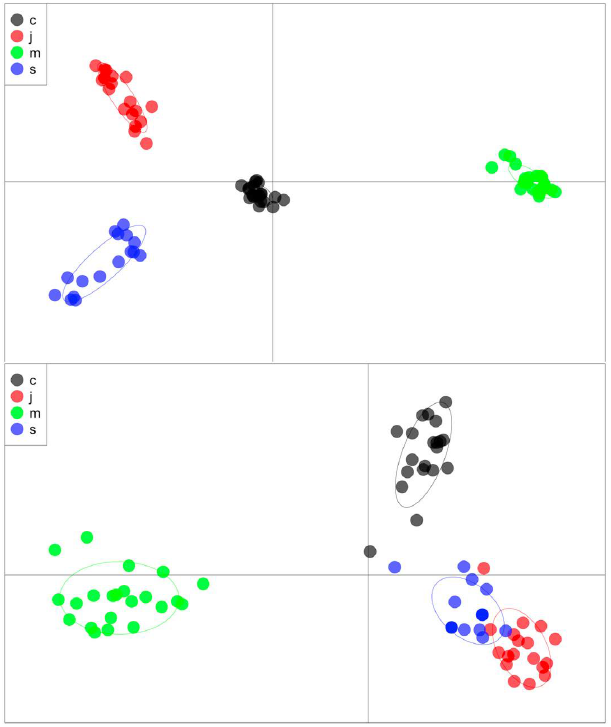
Discriminate Analysis of Principal Components (DAPC) using 3465 neutral SNPs (upper image) and 24 outliers SNPs (lower image). The first 4 PCs explained 29.8% of the variance (a-score = 0.621) when using neutral SNPs (upper image) and the first 5 PCs explained 82.8% of the variance (a-score = 0.594) when using outlier SNPs (lower image)

Allelic richness values (A_R_) ranged from 1.34 in populations J and S to 1.38 in population M. Observed heterozygosity (H_O_) values ranged from 0.361 in population S to 0.401 in population M, and were higher than expected heterozygosity (H_E_) in all populations suggestive of low inbreeding rates. Inbreeding coefficients (F_IS_) were negative in all populations indicating more heterozygous individuals than expected (Table 3). Effective population size values (Ne) were lowest for populations M and C (6.8 and 7.9, respectively; Table 3).

**TABLE 3.** Single nucleotide polymorphism (SNP) genetic diversity statistics by population. (N) number of individuals, (N_A_) number of SNP alleles, (H_O_) heterozygosity observed, (H_E_) heterozygosity expected, (A_R_) mean allelic richness, (F_IS_) inbreeding coefficient, and (N_e_) effective population size with 95% confidence intervals

|  | Population |  |  |  |
| --- | --- | --- | --- | --- |
|  | C | J | M | S |
| N | 20 | 18 | 22 | 16 |
| $N_A$ | 6888 | 6793 | 6900 | 6721 |
| $H_O$ | 0.388 | 0.370 | 0.401 | 0.361 |
| $H_E$ | 0.366 | 0.339 | 0.374 | 0.335 |
| $A_R$ | 1.37 | 1.34 | 1.38 | 1.34 |
| $F_{IS}$ | -0.050 | -0.073 | -0.060 | -0.065 |
| $N_e$ | 7.9<br>(7.8-7.9) | 23.4<br>(23.2-23.6) | 6.8<br>(6.8-6.8) | 11.4<br>(11.4-11.5) |

### 3.5 | COMPARISON OF POPULATION STRUCTURE USING MHC IA AND SNPS

For both MHC class IA and SNP data, STRUCTURE analyses identified an optimum of two clusters (*K* = 2), with individuals separating into clusters following similar patterns to fixation index (F_ST_) results (Table 2, Figure S8). MHC class IA STRUCTURE results indicated that 85% of individuals from population C grouped into one cluster, while only 36.8% individuals from populations M and S grouped into the same cluster. For SNP data, population M is the most divergent from the other three populations, which is in concordance with the DAPC results.

## 4 | DISCUSSION

Southern corroboree frogs exhibit phenotypic variation in *Bd* susceptibility which is associated with specific MHC class IA alleles and genome-wide SNPs. For example, MHC alleles Psco-UA*5, Psco-UA*9, and Psco-UA*23 were associated with either increased *Bd* infection loads or lower survival times. Multiple SNPs were suggestively correlated with *Bd* resistance including *RALGPS2*, which regulates immune cell proliferation and immunoglobulin Y (*IgY*), an antibody that binds and neutralizes pathogens. Despite dramatic recent declines, *P. corroboree* populations still contain sufficient standing genetic variation from which could be selected for improved survival. For example, detection of positive selection in the MHC class IA region of this species could suggest selection for *Bd* resistance. However, low interbreeding rates among closely interspersed *P. corroboree* populations confirm natural history observations for this species of low vagility. This factor explains declines in *P. corroboree* despite evidence for sufficient additive genetic *Bd* resistance in the species. Therefore, *P. corroboree* may benefit from genetic manipulation to improve *Bd* resistance in order to overcome natural history constraints that prevent optimal selection conditions.

### 4.1 | PHENOTYPIC DIFFERENCES AMONG POPULATIONS

*P. corroboree* exhibit phenotypic variation in resistance to *Bd* infection at the population level. This was evident in both the days survived and infection load through time (Figure 1). One population (M) was distinct from the other three populations by having the longest survival, lowest infection loads, and greatest amount of individuals that survived until the end of the experiment.

### 4.2 | GENOTYPE-PHENOTYPE ASSOCIATIONS

Phenotypic differences in *Bd* resistance were associated with genetic variance of the MHC and genome-wide SNPs. Three MHC alleles were associated with increased *Bd* susceptibility in individual frogs. Of these, Psco-UA*5 was least common in the more *Bd* resistant M population. MHC supertypes were not associated with resistance, however, supertype ST8 was associated with increased susceptibility. Although this supertype was relatively common, it was more common in the frogs that died (86%) than in survivors (40%), and least common in the more resistant population M (Table S4, Figure 3a).

Our GWAS did not identify any SNPs that were significantly associated with *Bd* resistance, which may be the result of the limited sample size available for this study combined with the potential polygenic nature of the traits analyzed here. One SNP (173) was suggestively negatively associated with days survived (Figure 4). This SNP has closest homology to the *RALGPS2* gene of *X. tropicalis* that encodes a guanine nucleotide exchange factor (GEF) for the GTPase RALA (Tables 4 and S2). These molecular switches are involved in multiple cell processes such as cell differentiation and proliferation (Alberts *et al.* 2002) and may potentially influence *Bd* immunity by regulating epidermal or immune cell proliferation, which are important predictors of *Bd*-related mortality (Ellison *et al.* 2014a). Several of the top 10 SNPs associated with maximum infection load (e.g., 1894, 1895) have sequence homology to an alpha-L-tissue fucosidase (*FUCA1*). This enzyme cleaves fucose containing glycoproteins, and is involved in the immunoregulation of leukocyte migration during inflammation (Ali *et al.* 2008). Significantly, poor immune cell recruitment is observed in the skin of *Bd-*infected frogs, due to pathogen-produced immunosuppressants that cause apoptosis (Fites *et al.* 2013). It remains to be seen whether suppression of inflammation in chytridiomycosis involves immunoregulation via fucosidases, although preliminary support indicates the increased transcription of fucose binding lectin in frogs after exposure to *Bd* (Ribas *et al.* 2009). A SNP with homology to immunoglobulin Y (IgY) was also identified as a top SNP in this study (3440). IgY is functionally analogous to mammalian IgG, which is essential for pathogen recognition and control (Warr *et al.* 1995). The contribution of IgY to *Bd* immunity varies across species. *Xenopus laevis* immunized against *Bd* produced a strong pathogen-specific IgY response (Ramsey *et al.* 2010), whereas IgY response was suppressed or unaffected in other species (Fernández-Loras *et al.* 2017; Poorten *et al.* 2016; Young *et al.* 2014).

**TABLE 4.**
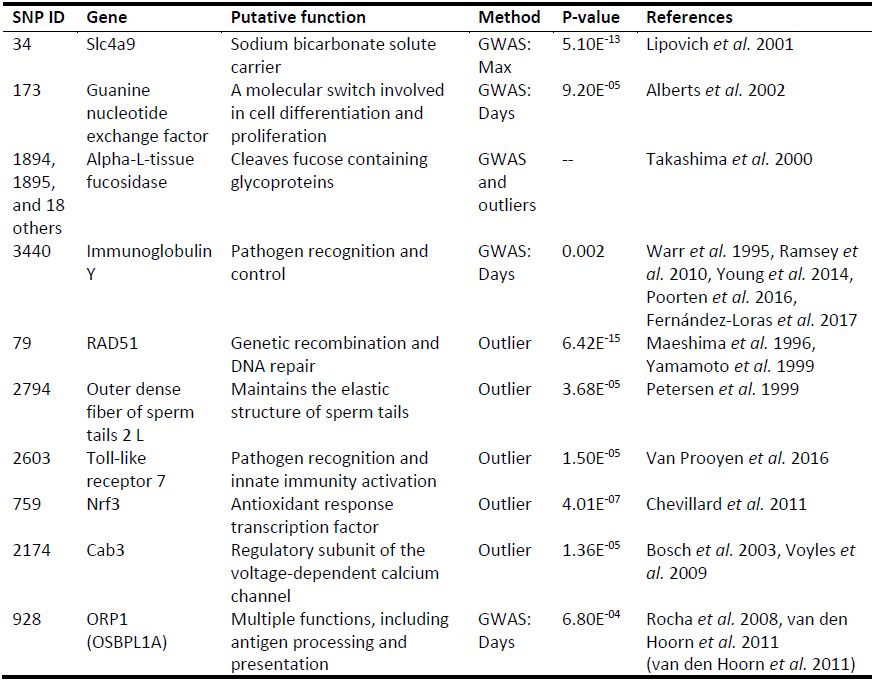
Putative gene functions of notable SNP loci identified by GWAS and outlier analyses

### 4.3 | GWAS LIMITATIONS

Although GWAS is a powerful tool for detecting SNP-phenotype associations, this method is limited in its ability to detect SNPs with low to moderate effect sizes (Harrisson *et al.* 2014). For example, Barson *et al.* (2015) was successful at identifying a large effect locus controlling age of maturity in Atlantic salmon, but other studies have failed to detect significant loci (e.g., Santure *et al.* 2015). In many cases where significant variants have been associated with phenotypic traits, they only explain a relatively small proportion of the variance (e.g., Bérénos *et al.* 2015; Silva *et al.* 2017). This suggests a quantitative genetic architecture, and that causal loci are being overlooked due to a lack of power to detect multiple small effects (Maher 2008; Yang *et al.* 2010).

Therefore, GWAS performs best with large sample sizes and high SNP coverage (Visscher *et al.* 2017), making GWAS investigations of threatened or non-model species challenging. For *P. corroboree*, power calculations based on the average predicted genomic relationships from the individuals used in this study indicate that to achieve 99.7% power, a sample size of (n = 1000) is required if narrow sense heritability of *Bd* resistance is (h^2^) > 0 (see Methods S2 for description of power calculations). Future studies with *P. corroboree* should increase the sample size in order to improve GWAS power.

### 4.4 | POPULATION GENETIC STRUCTURE

Southern corroboree frog populations showed significant evidence of genetic structure between all four populations studied, with population M being the most differentiated in pairwise comparisons across all datasets, and populations J and S the least differentiated (Table 2, Figures 5 and S8). The population divergence estimates for this species are consistent with the life history information (i.e., high site fidelity, low vagility; Hunter 2000), and microsatellite data indicates that interbreeding among populations separated by even a few km is low (Morgan *et al.* 2008). These factors have likely contributed to the demise of *P. corroboree* after the introduction of *Bd*, despite evidence of *Bd* resistance within some individuals. This is because *Bd* resistance genes are unlikely to spread across the landscape due to low interbreeding rates.

Populations C and M showed low effective population sizes (Ne), which reflects the recent demographic history of this species in that it went from highly abundant to functionally extinct within the last ~30 years (Hunter *et al.* 2010b; Morgan *et al.* 2008; Osborne *et al.* 1999). Although we observed an excess of heterozygotes (H_O_) and low inbreeding rates (F_IS_)—suggestive of a large, diverse population—this may be due to the relatively low sample sizes available for this study and/or the effects of the recent drastic reduction in population size that might not yet have impacted inbreeding estimates.

### 4.5 | EVOLUTION OF THE MHC CLASS IA AND SNPS

Tests of selection indicate that codons of the putative MHC peptide binding region pockets are under positive selection in *P. corroboree*, suggesting that these amino acid residues provide a survival advantage to *Bd* infected hosts.

Even though we did not detect any MHC alleles associated with *Bd* resistance, our evidence that three MHC alleles and one supertype are associated with higher susceptibility supports the hypothesis that *P. corroboree* MHC has a functional role in *Bd* immunity. In other frog species, MHC class IIB alleles and supertypes have been correlated with increased *Bd* resistance (e.g., Bataille *et al.* 2015; Savage & Zamudio 2011) likely due to higher binding affinity for *Bd* peptides. This correlation is also probable for MHC class IA, as has been demonstrated in pathogen systems of humans and other species (Aguilar *et al.* 2016; International H. I. V. Controllers Study *et al.* 2010; Koch *et al.* 2007; Madsen & Ujvari 2006). Future investigations should develop locus specific primers or apply next-generation sequencing to improve the confidence level of genotyping (see Galan *et al.* 2010). Another approach would be to knock-in putative *Bd* immunity-associated MHC alleles using gene editing technologies, such as CRISPR-Cas9 (Doudna & Charpentier 2014), and measure gene effects directly via *Bd* challenge experiments.

Of the 24 SNPs that were population outliers (Tables 4 and S11), there are several that likely play an important role in *Bd* immunity due to their homology to pathogen response genes in other species. One of the SNPs under positive selection is a *RAD51* homolog (79). In *Xenopus*, this gene is involved in genetic recombination and DNA repair, and is likely involved in meiotic recombination due to high expression levels in testes and ovaries (Maeshima *et al.* 1996). *RAD51* has also been shown to be strongly expressed in newt testes during spermatogenesis (Yamamoto *et al.* 1999), a process that is increased in *Bd*-infected frogs (Brannelly *et al.* 2016a). Interestingly, another SNP identified under selection in our study had homology to an outer dense fiber that maintains the elastic structure of sperm tails (2794), further highlighting the potential role of increased reproductive effort as a response to *Bd* infection.

An additional SNP outlier (2603) has homology to Toll-like receptor 7 (TLR7), which is involved in pathogen recognition and activation of innate immunity via production of specific cytokines. In response to the intracellular fungal pathogen *Histoplasma capsulatum*, murine dendritic cells require TLR7 to control fungal growth and activate T cells via interferon-γ (Van Prooyen *et al.* 2016). It is possible that this gene may also be involved in the response of frogs to *Bd* infection. Other relevant SNPs outliers include those that may act in response to *Bd*-induced effects on hematopoietic tissue (Brannelly *et al.* 2016b), and electrolyte transport and cardiac function (Voyles *et al.* 2009). For example, the SNP *nrf3* (759), is an antioxidant response transcription factor with a protective role in hematopoietic tissues (Chevillard *et al.* 2011); while the SNP, *cab3* (2174), is a regulatory subunit of the voltage-dependent calcium channel that is downregulated in rabbit models with rapid heart rates (Bosch *et al.* 2003). Regulation of calcium channels may impact chytridiomycosis outcomes since *Bd* kills hosts by interfering with electrolyte homeostasis causing eventual cardiac arrest (Voyles *et al.* 2009).

### 4.6 | IMPLICATIONS FOR THE CAPTIVE BREEDING PROGRAM

Although *P. corroboree* are functionally extinct in the wild, our results suggest that the captive populations (i.e., in terms of allelic richness and heterozygosity) still host good levels of the genetic diversity which may constitute a potential genetic input to wild populations. The immunogenetic variation for *Bd* resistance suggests that genetic manipulation methods could be used to increase species-wide *Bd* resistance to ensure the successful reintroduction of *P. corroboree.* The IUCN’s Amphibian Conservation Action Plan recommends establishing captive assurance colonies for amphibians threatened by *Bd* (Gascon *et al.* 2007; Wren *et al.* 2015). In response to these recommendations, the amphibian ARK (AARK) was setup to manage and advise captive breeding efforts, and this successful program currently consists of 188 projects worldwide (AARK database 2017). However, because *Bd* cannot be extirpated from the environment, reintroduction projects are unlikely to be effective unless animals with increased resistance are released. The best approaches for increasing disease resistance in captive breeding programs come from livestock and agriculture where they have been successfully applied for over 100 years (Hickey *et al.* 2017). Methods that utilize multiple genetic markers, such as genomic selection, are ideal for increasing disease resistance for wildlife because they have the highest genetic gain (i.e., change in mean trait per year) and the lowest inbreeding rates (Daetwyler *et al.* 2007; Hickey *et al.* 2017; Meuwissen *et al.* 2016). However, before genetic manipulation methods can be applied to corroboree frogs and other species to be released into the wild, several precautions should be undertaken. Most importantly, SNPs used for increasing resistance should be determined from robust, well-powered studies so that conservation managers can be confident of their impact. Additionally, genetically modified animals should be trialed in the field across the range of their environment to ensure that they do not have reduced overall fitness in the wild.

## 5 | CONCLUDING REMARKS AND FUTURE DIRECTIONS

Southern corroboree frog populations have phenotypic and genetic variation in *Bd* susceptibility. Hence, they have the potential to be genetically manipulated to increase *Bd* resistance. We also show that despite functional extinction in the wild, there is still substantial genetic variation in this species within the captive assurance collection. This pilot study is a first step towards using genomic approaches to investigate polygenic immunity to *Bd*. Future studies should further examine the role that these identified SNPs and MHC variants play in *Bd* resistance. This could be investigated by genetically engineering frogs with gene knock-in approaches or applying genomic selection to increase the frequency of the genes of interest, followed by *Bd*-challenge experiments to measure their impact on the resistance phenotype. Additional studies should also strive to fully characterize the genetic architecture and heritability of *Bd* resistance by performing high resolution QTL association mapping and high-powered GWAS. Lastly, high quality genomic resources for amphibians are required to inform GWAS and comparative genome analyses.

## ACKNOWLEDGMENTS

We extend sincere thanks to Gerry Marantelli for providing the corroboree frogs used in this study. We are grateful to Collin Storlie, Rhondda Jones, Donald McKnight, and the JCU eResearch team for providing R scripting and statistical assistance, Rebecca Webb, Jennifer Hawkes, Ket Fossen, and Cam De Jong for assistance with animal husbandry, Sara Bell for disease testing assistance, David Hunter for conservation agency support, Michael McFadden, Peter Harlow, and Raelene Hobbs for animal husbandry advice and Kyall Zenger, John Eimes and Arild Husby for discussion of analysis approaches. Funding was provided by the Australian Research Council grants LP110200240 and FT100100375, New South Wales Office of Environment and Heritage, Taronga Zoo, experiment.com crowdfunding grant “Can we stop amphibian extinction by increasing immunity to the frog chytrid fungus”, and Queensland Department of Science, Information Technology and Innovation Accelerate Fellowship grant 14-218.

## DATA ACCESSIBILITY

MHC class IA DNA sequence data is available on GenBank (acc#’s xx-xx). The data generated from this study is accessible on Dryad xxx.

## AUTHOR CONTRIBUTIONS

TAK and CNSS drafted the manuscript. TAK conducted MHC sequencing. TAK and QL conducted MHC analysis. CNSS and TAK performed association and population genetic analyses. LAB performed infection study and *Bd* quantitation. All authors contributed to study design, writing the manuscript, and approved the final version.

## SUPPORTING INFORMATION

Additional supporting information may be found in the online version of this article

**Table S1** MHC class IA Sanger sequencing information

**Table S2** MHC class IA genotypes

**Table S3** MHC class IA allele frequencies

**Table S4** MHC class IA supertype (ST) frequencies

**Table S5** Results of GLS for MHC class IA alleles associated maximum infection load

**Table S6** Results of GLS for MHC class IA alleles associated with days survived

**Table S7** Results of GLS for MHC class IA supertypes associated with maximum infection load

**Table S8** Results of GLS for MHC class IA supertypes associated with days survived

**Table S9** Descriptive information on the top 10 SNPs for the GWAS models

**Table S10** BLAST results of the top 10 SNPs for the GWAS models

**Table S11** BLAST results for SNP population outliers

**Figure S1** Body condition by population of frogs from *Bd* treatment group

**Figure S2** Comparison of ω across the *P. corroboree* MHC class IA amino acid alignment

**Figure S3** Discriminate Analysis of Principal Components (DAPC) of MHC class IA supertypes

**Figure S4** Evolutionary relationships of MHC Class IA nucleotide sequences and MHC supertypes

**Figure S5** Genetic diversity of MHC class IA in *P. corroboree* populations

**Figure S6** Influence of supertype 8 on survival

**Figure S7** Manhattan plots with −log10 p-values from the GWAS association

**Document S1** The detailed description of the methods

